# Mapping temperature-sensitive mutations at a genome-scale to engineer growth-switches in *E. coli*

**DOI:** 10.1101/2023.06.01.543195

**Authors:** Thorben Schramm, Vanessa Pahl, Hannes Link

## Abstract

Temperature-sensitive (TS) mutants are a unique tool to perturb and engineer cellular systems. Here, we constructed a CRISPR library with 15,120 *Escherichia coli* mutants, each with a single amino acid change in one of 346 essential proteins. 1,269 of these mutants showed temperature-sensitive growth in a time-resolved competition assay. We reconstructed 94 TS mutants and measured their metabolism under growth arrest at 42°C using metabolomics. Metabolome changes were strong and mutant-specific, showing that metabolism of non-growing *E. coli* is perturbation-dependent. For example, 24 TS mutants of metabolic enzymes overproduced the direct substrate-metabolite due to a bottleneck in their associated pathway. A strain with TS homoserine kinase (ThrB^F267D^) produced homoserine for 24 hours, and production was tunable by temperature. Finally, we used a TS subunit of DNA polymerase III (DnaX^L289Q^) to decouple growth from arginine overproduction in engineered *E. coli*. These results provide a strategy to identify TS mutants *en masse* and demonstrate their large potential to produce bacterial metabolites with non-growing cells.

## Introduction

Targeted perturbations of cellular networks are key to understand and engineer their function. Gene deletions, for instance, can improve microbial production strains (Burgard *et al*, 2003), and knockout mutant libraries enabled systematic analyses of gene-gene networks (Tong *et al*, 2004), gene-metabolite networks (Fuhrer *et al*, 2017; Mülleder *et al*, 2016), and gene-regulatory networks (Kemmeren *et al*, 2014). However, gene deletions are static and irreversible perturbations, and they are not feasible if the gene of interest is essential for cell growth. RNA interference (Na *et al*, 2013) or CRISPR interference (Qi *et al*, 2013) allow inducible knockdowns of essential genes, and these methods were used to construct synthetic regulatory circuits (Santos-Moreno *et al*, 2020; Qi *et al*, 2013) and dynamic growth switches (Li *et al*, 2016).

Temperature-sensitive (TS) mutations are an alternative method to perturb essential genes. At low (permissive) temperatures, genes with a TS mutation encode a functional product, while at higher (non-permissive) temperatures, the gene product is not functional. Several molecular mechanisms can lead to thermal sensitivity, the most important of which induce changes in protein stability or changes in protein folding. Thermolabile mutant proteins, for instance, unfold at higher temperatures, which reduces their activity or inactivates the protein completely. In contrast, proteins with temperature-sensitive folding are not correctly folded at higher temperatures (Haase-Pettingell & King, 1997; Sadler & Novick, 1965). Other TS mechanisms alter interactions of the TS protein with other molecules, such as a TS mutant of the Drosophila muscle regulator Mef2, which changes DNA binding upon temperature changes (Lovato *et al*, 2009).

TS mutations provide several unique advantages compared to other perturbation methods like RNA or CRISPR interference. First, a single mutation is often sufficient for a TS phenotype and, thus, auxiliary components like dCas9 (Qi *et al*, 2013) or small regulatory RNAs (Na *et al*, 2013) are not required. Second, many TS mutations enable fast perturbations, especially if the mutant protein is thermolabile and unfolds within seconds or minutes (Plaza del Pino *et al*, 2000). Third, TS mutations have almost no polar effects since mutations are small and will mainly affect a single gene product. Finally, temperature shifts are extremely versatile, well-controllable, and reversible. Focused ultrasound, for instance, can control TS proteins *in vivo* (Piraner *et al*, 2017), and almost all bioreactors are equipped with temperature control.

TS mutants were used to engineer diverse cellular systems with applications in medical and industrial biotechnology (Kasari *et al*, 2022; Weber, 2003; Harder *et al*, 2018; Lynch *et al*, 2019, 2016; Cho *et al*, 2012; Wang *et al*, 2021; Piraner *et al*, 2017; Schramm *et al*, 2020). However, most applications are based on known temperature-sensitive mutations, such as the transcriptional repressor CI857 from *Escherichia virus Lambda* (Kasari *et al*, 2022; Wang *et al*, 2021; Harder *et al*, 2018). The search for new TS mutants was mainly driven by large-scale genetic screens in yeast (Costanzo *et al*, 2016). These efforts required a comprehensive collection of TS mutants of *Saccharomyces cerevisiae*, which, for the most part, were constructed with random mutagenesis approaches (Ben-Aroya *et al*, 2008; Kofoed *et al*, 2015; Li *et al*, 2011).

Currently, there is no comprehensive collection of TS *Escherichia coli* strains. The *Coli* Genetic Stock Center Database (Berlyn, 1999) lists 178 temperature-sensitive mutations in 116 protein-coding genes. We found an additional 41 temperature-sensitive *E. coli* strains in the literature that cover another 15 genes (Suppl Table 1). To our knowledge, 87 TS *E. coli* strains with mutations in 32 genes were whole genome sequenced. All the 87 strains had at least one amino acid substitution, and 29 had more than one. These temperature-sensitive *E. coli* strains were relevant to study the function of individual genes and contributed to important findings like how DNA is replicated (Blinkova *et al*, 1993; Georgescu *et al*, 2008; Hansen & Atlung, 2018; Saluja & Godson, 1995; Vandewiele *et al*, 2002).

Here, we used a high-throughput approach to construct and identify temperature-sensitive mutants. We used a CRISPR method to construct 15,120 *E. coli* strains, each with a single amino acid change in one of 346 essential proteins, and we measured their growth at two temperatures. Based on these results, we constructed a panel of 94 TS *E. coli* strains with single mutations and analyzed their growth and metabolism by metabolomics. Many TS mutants of enzymes accumulated the direct substrate-metabolite. For example, TS variants of homoserine O-succinyltransferase (MetA^F285W^) and homoserine kinase (ThrB^F267D^) overproduced homoserine, and the production was tunable by temperature. Finally, we used a TS subunit of DNA polymerase (DnaX^L289Q^) to control growth of an arginine overproducing *E. coli*.

## Results

### 15,120 E. coli mutants with single amino acid changes in 346 essential proteins

We used a modified version of a CRISPR-Cas9 method (Garst *et al*, 2017) to create a library of 15,120 *E. coli* strains, each with a different amino acid change in an essential gene. As a starting point, we selected 352 proteins that are essential for growth of *E. coli* on minimal glucose medium (Goodall *et al*, 2018; Patrick *et al*, 2007). We then designed single amino acid changes that may cause temperature-sensitivity of the respective protein using the TSpred algorithm (Tan *et al*, 2014; Varadarajan *et al*, 1996). To reduce the design space, we included only amino acid changes to alanine, aspartate, glutamine, proline, and tryptophan (Tan *et al*, 2014; Varadarajan *et al*, 1996). Per protein, we designed up to 50 amino acid changes (10 sites each with 5 substitutions) considering the following design rules: (1) a minimal distance between the protospacer adjacent motif (PAM) site and the mutation site to maximize editing efficiency, (2) a maximal number of possible amino acid changes at a given site, and (3) minimal CRISPR off-targets of the single guide RNA (sgRNA).

For 154 genes, we found less than 50 amino acid changes either because the number of predicted sites were limited or due to constraints by the design rules. For example, TSpred predicted no temperature-sensitive mutations in *rpmA* and *rpmH,* which encode small ribosomal proteins with 85 and 45 amino acids, respectively. Another set of genes (*leuL*, *rplU*, *rpmC*, and *rpsI*) had no substitutions that fulfilled our design rules. In total, we designed 16,038 single amino acid changes and inserted the respective mutations into 346 essential genes from various functional categories (Suppl Table 2).

The CRISPR-based genome editing method uses homologous recombination by *Escherichia virus Lambda* Red (Garst *et al*, 2017). Templates for homologous recombination were 85 bp-long DNA sequences that had the desired mutation to introduce single amino acid changes and an additional silent mutation at the protospacer adjacent site. The homologous DNA sequence was encoded on a plasmid next to the respective sgRNA and functioned as a strain-specific barcode. We constructed the plasmids in a pooled approach using 200 bp array-synthesized oligonucleotides and measured the library composition directly after construction by deep sequencing (Fig 1A and Fig EV 1). Out of all 16,038 designed amino acid substitutions, 15,582 (97%) were present in the plasmid library.

**Figure 1.**
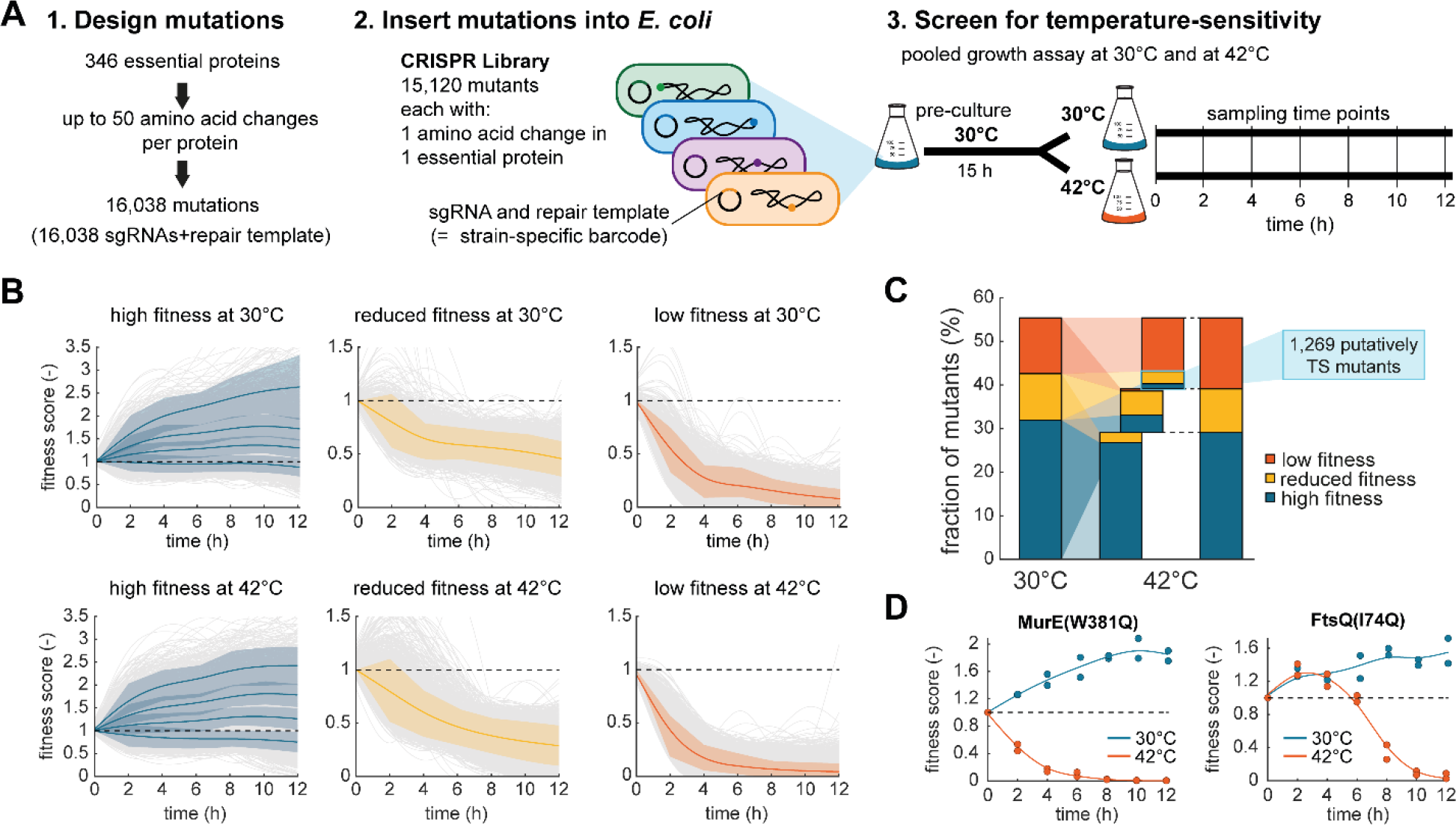
A CRISPR screen with 15,120 *E. coli* mutants identifies temperature-sensitive mutations. **A,** Schematic of the CRISPR screen. 16,038 sgRNAs plus repair templates (barcodes) were designed to introduce amino acid changes in 346 essential proteins (step 1). 15,120 of the barcodes were present in the final CRISPR library (step 2). The CRISPR library was cultured at 30°C and at 42°C (n = 2 replicates). Strain-specific barcodes (sgRNA and repair template) were sequenced every 2 h to determine the composition of the library (step 3). **B,** K-means clustering of fitness scores of 8,884 strains in the CRISPR library. Time-course data were clustered into k = 6 clusters per temperature. The fitness scores were calculated by normalizing the read counts of the barcode of each mutant to the total number of reads per sample and to the first time point. Grey curves are the moving average of the mean of n=2 replicates. Colored lines are cluster means and shaded areas their standard deviation (blue: 4 cluster with high fitness, yellow: 1 cluster with reduced fitness, red: 1 cluster with low fitness). **C,** Relative composition of the CRISPR library at 30°C and 42°C. Blue indicates high fitness, yellow a reduced fitness, and red low fitness. The bar graph in the middle connects the 30°C and 42°C data. The light blue box indicates putative TS mutants. **D,** Examples of fitness score dynamics of two strains that show temperature sensitivity (MurE^W381Q^ and FtsQ^I74Q^). Dots show data from n = 2 replicates per temperature. The lines are the moving average through the means. Blue: 30°C culture. Red: 42°C culture.

Next, we used the plasmid library for the transformation of an *E. coli* strain, which carried a second plasmid with Cas9 and the *Lambda* Red system (Suppl Fig 1). In these strains, we induced Cas9 expression and *Lambda* Red-mediated recombination to obtain the final CRISPR library. This library contained 15,120 of all designed 16,038 single amino acid substitutions (94%) and targeted all 346 genes that we included in the initial library design.

### Time-resolved competition assays identify putative TS mutants

After constructing a CRISPR library with 15,120 mutants, we sought to identify mutants that are temperature-sensitive. For this purpose, we used a time-resolved competition assay, in which we cultivated the pooled CRISPR library at 30°C and 42°C (Fig 1A). First, we grew the CRISPR library for 15 h on minimal glucose medium at 30°C and expected that strains with a strong growth defect would disappear from the library during this pre-culture phase. After 15 h, the pre-culture was then used to inoculate two main cultures: a 30°C culture and a 42°C culture. These two main cultures were incubated for 12 h. Every 3 h, the cultures were back diluted into fresh medium to avoid limitations of oxygen and nutrients. Every 2 h, we determined the composition of the library by deep sequencing of the strain-specific barcodes, which was reproducible between two independent experiments (Suppl Fig 2). Fitness scores of single mutants were determined by normalizing the read counts of their barcodes to the total number of reads and the first time point.

Out of all 15,120 strains, 6,236 dropped out from the library after the 15-hour pre-culture phase (strains with average reads < 15), presumably because their mutation inactivated the protein, and we discarded these strains from further analysis. For the remaining 8,884 strains, we explored dynamic patterns in the main cultures with k-means clustering (Fig 1B). This analysis revealed that 5,118 strains had a high fitness at 30°C, 1,712 strains had mild fitness defects at 30°C, and 2,054 strains had strong fitness defects at 30°C. Most strains were not affected by temperature. This means that 96% of the strains with a fitness defect at 30°C also had a fitness defect at 42°C (Fig 1C). Similarly, 84% of the mutants with a high fitness at 30°C also had a high fitness at 42°C. However, 1,269 mutants (8.4% of the library) had a higher fitness at 30°C than at 42°C, thus indicating that these strains are TS mutants (blue box in Fig 1C). The fitness scores of the putative TS mutants showed a wide range of dynamics. Some TS mutants like MurE^W381Q^ grew well at 30°C, and disappeared fast from the library in the 42°C culture (Fig 1D). Other mutants, like FtsQ^I74Q^, disappeared with a time-delay from the 42°C culture (Fig 1D).

In summary, we constructed a CRISPR library with 15,120 strains and identified 1,269 strains that are putative TS mutants. Next, we reconstructed some of these mutants and examined them individually.

### A panel of 94 TS mutants shows distinct growth-temperature relationships

To reconstruct the best TS mutants, we scored temperature-sensitivity of all strains in our library by a set of criteria (Suppl Fig 3). With this approach, we obtained high-scoring TS mutations for 250 essential genes in the CRISPR library (Suppl Table 3) and constructed a sub-library with these 250 strains using the same CRISPR-Cas9 method (Garst *et al*, 2017). Sequencing of the sub-library showed that all 250 strains were present in the library, but their relative abundance varied markedly between 0.0007% and 2.4% (Fig 2A). If these differences in the abundance of single strains were caused by experimental variation, we expected that cloning the sub-library a second time would reduce the variation. However, the relative abundance of the 250 strains remained remarkably constant between the two cloning rounds (Fig 1A), thus indicating that mutant-specific parameters (e. g., editing efficiency) influenced the abundance of single strains in the library and not experimental variation during cloning.

**Figure 2.**
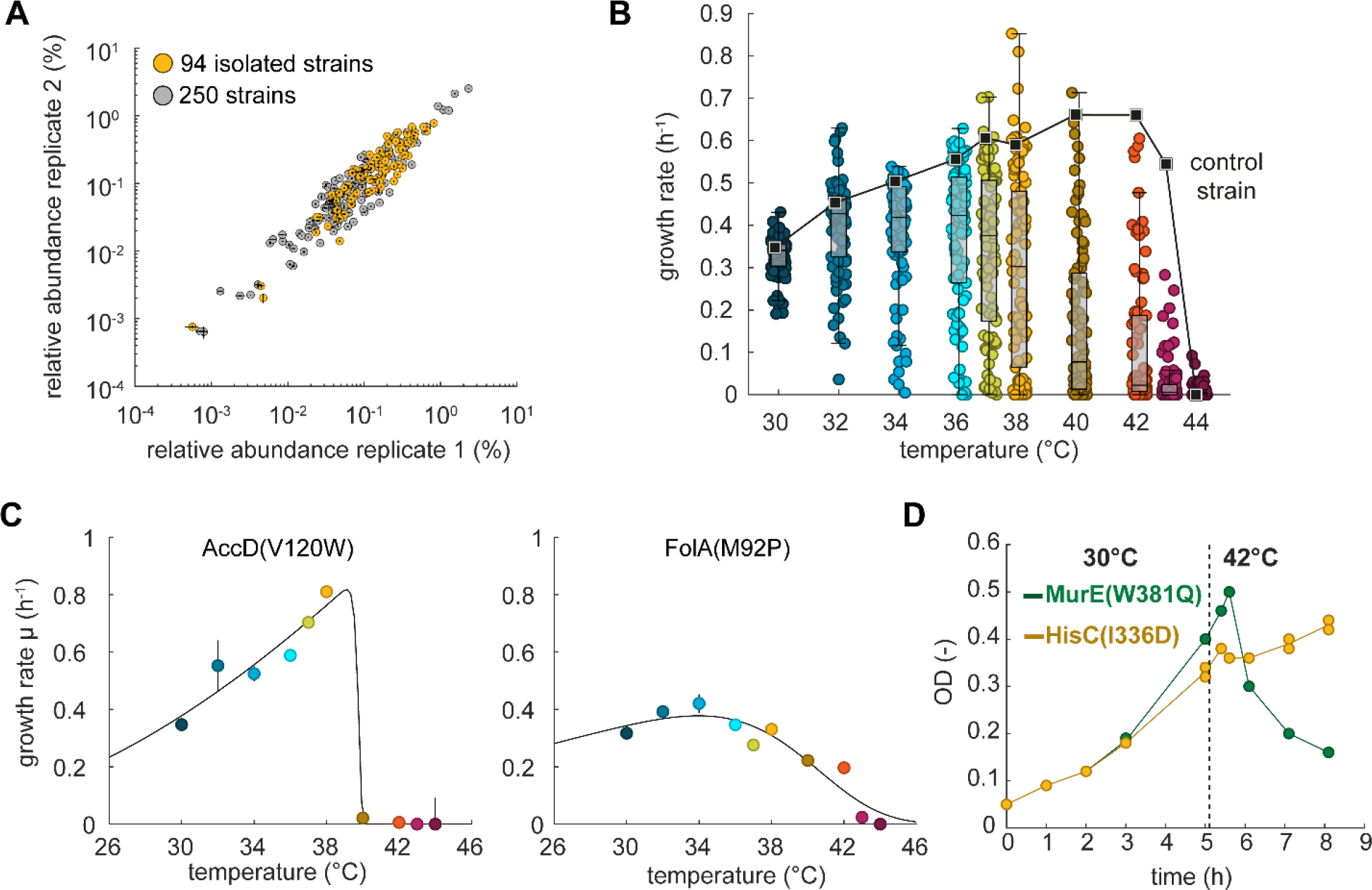
**Reconstruction of 94 TS mutants and characterization of growth-temperature relationships.** **A,** Relative abundance of strains in the pooled sub-library of 250 putative TS mutants. The strain library was constructed twice (replicate 1 and 2), and dots are the mean of n= 2 technical sequencing replicates. The black lines indicate the difference between the two technical replicates. The read counts of single strains were normalized to the total number of reads. Yellow dots indicate 94 TS mutants that were isolated from the pooled sub-library. **B,** Maximal specific growth rates of 94 TS mutants and a control strain at ten different temperatures between 30°C and 44°C. Dots are the mean (n = 3). Black squares and the line indicate the control strain. The box whisker plots show the median and 25^th^/75^th^ percentiles. **C,** Examples of maximal specific growth rates of the TS mutants AccD^V120W^ and FolA^M92P^ at different temperatures (Fig EV 2 shows all 94 TS mutants). Dots are the mean, vertical black lines indicate the standard deviation (n = 3). The black line was calculated by fitting an Arrhenius-type function to the data (see also Suppl Fig 4). **D,** Growth dynamics of the MurE^W381Q^ strain (green) and the HisC^I336D^ strain (yellow) during a temperature shift from 30°C to 42°C. Dots show data from n = 2 replicates, and the lines connect the mean.

We then isolated random strains from the sub-library and tested them for temperature-sensitivity until we had a panel of 94 unique TS mutants. The 94 TS mutants covered genes from all functional categories, except ribosomal subunits. To characterize growth-temperature relationships of the 94 TS mutants, we cultured them together with a non-edited control strain at 10 different temperatures between 30°C and 44°C (Fig 2B, Fig EV 2, and Suppl Table 4). At 30°C, most TS mutants had similar specific growth rates as the control strain (0.35 h^-1^), whereas specific growth rates varied markedly at higher temperatures. At 38°C, for example, specific growth rates of the 94 mutants varied between no growth and 0.85 h^-1^, which is close to the maximal specific growth rate that is possible for *E. coli* on minimal glucose medium (Monk *et al*, 2016). The non-edited control strain had a higher specific growth rate at 42°C (0.66 h^-1^) than at 37°C (0.61 h^-1^), which is consistent with previous studies (Schmidt *et al*, 2016). At 43°C, the specific growth rate of the control strain decreased slightly to 0.54 h^-1^, whereas all TS mutants showed a strong growth defect at this temperature.

The growth-temperature relationships of all TS mutants and the control strain followed an empirical Arrhenius-type function with an activating term and an inactivating term (Suppl Fig 4A/B, median R^2^ value = 0.96). These two terms may reflect that higher temperatures activate overall metabolism, but, at the same time, they inactivate the TS protein. Although all growth-temperature relationships qualitatively matched the Arrhenius-type function, they differed quantitatively across the 94 TS mutants, resulting in distinct parameters (Fig EV 2). Further clustering separated strains with a switch-like response from those with a gradual response (Suppl Fig 5). 52 TS mutants and the control strain had a switch-like response to temperature increases, which means that, at some point, a small temperature change led to large changes in the growth rate. For example, the AccD^V120W^ mutant switched from 0.81 h^-1^ to 0.02 h^-1^ between 38°C and 40°C (Fig 2C). Another set of 42 mutants showed a more gradual decrease of the growth rate in response to temperature increases. The FolA^M29P^ mutant, for instance, required a temperature increase of 9°C to switch from its maximal growth rate to no growth (Fig 2C).

Next, we wondered how fast a TS mutant switches from growth to no growth upon temperature increases. Therefore, we selected two TS mutants from our panel of 94 strains that showed a fast response in the pooled competition assay: the MurE^W381Q^ mutant (Fig 1D) and the HisC^I336D^ mutant (Suppl Fig 6). At 30°C, the two strains grew normally, and, after 5 h, we transferred them to 42°C (Fig 2D). After this temperature shift, the MurE^W381Q^ mutant was able to grow for another 30 min, but then the optical density of the culture decreased, indicating cell lysis. Perturbation of the MurE reaction may have caused a limitation in peptidoglycan biosynthesis, and it is thus possible that cell lysis of MurE^W381Q^ at 42°C resembles the bactericidal effect of antibiotics that target peptidoglycan metabolism (Bush & Bradford, 2016; Williams & Bardsley, 1999). The HisC^I336D^ strain also showed a reduction of growth after 30 min at 42°C, but did not lyse like the MurE^W381Q^ strain. Since the histidinol-phosphate aminotransferase (HisC) catalyzes the seventh step in histidine synthesis (Grisolia *et al*, 1985), the growth arrest of the HisC^I336D^ mutant at 42°C was probably due to a histidine limitation.

In conclusion, we constructed a panel of 94 TS mutants that had growth defects at higher temperatures but grew similar as an unedited control strain at 30°C. Although the growth-temperature relationships differed across the 94 mutants, they all followed an empirical Arrhenius-type function. Dynamic temperature shifts demonstrated that growth of the TS mutants MurE^W381Q^ and HisC^I336D^ responded within 30 min after a shift from 30°C to 42°C, either with cell lysis (MurE^W381Q^) or with a growth arrest (HisC^I336D^). Thus, different TS mutants have distinct growth phenotypes, and we wondered if these differences occur also at the level of metabolism.

### TS mutants show distinct metabolomes and increases of substrate metabolites

To probe how the TS mutations affect cellular metabolism, we analyzed the metabolomes of the 94 TS mutants (Fig 3A). For this purpose, we cultured the TS mutants and an unedited control strain at 42°C and measured metabolite level after 16 h by flow-injection time-of-flight mass spectrometry (FI-MS) (Fuhrer *et al*, 2011; Farke *et al*, 2023). FI-MS detected 325 metabolites, of which 219 showed strong increases in at least one TS mutant (mod. z-score > 3). We analyzed if these strong metabolome changes were a global response to the growth reduction at 42°C or if the metabolome responses were specific to the TS mutants. Most metabolite increases were specific, because 53 metabolites increased in a single TS mutant and only 14 metabolites increased in more than 10 mutants (increases with mod. z-score > 3).

**Figure 3.**
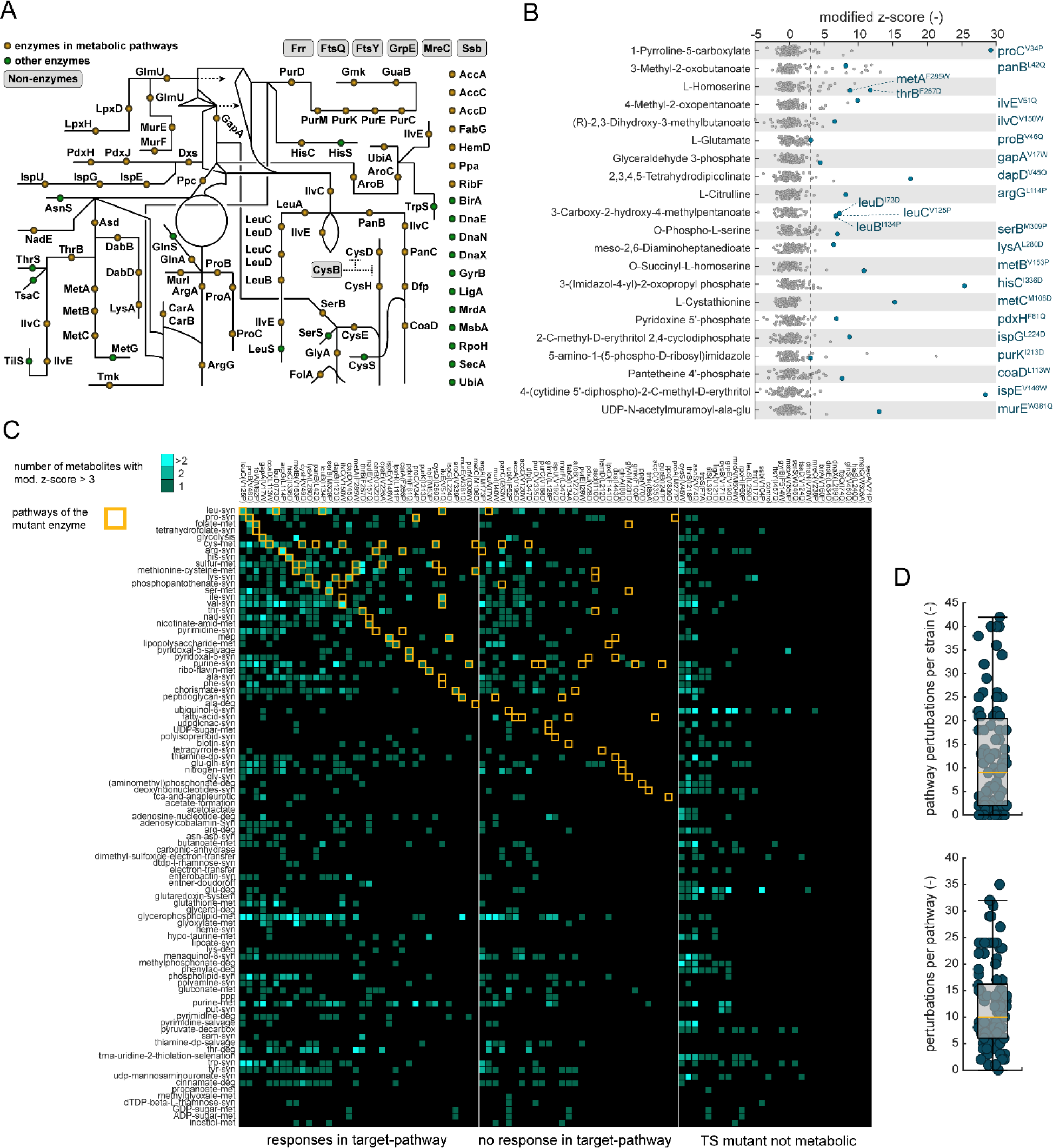
**Metabolome responses of 94 TS mutants are strong, mutant-specific, and metabolism-wide.** **A,** Location of TS mutants in the metabolic network. 66 TS mutants affect metabolic enzymes (yellow dots). 22 TS mutants are non-metabolic enzymes (green dots). 7 TS mutants affect proteins without enzymatic function (grey boxes). **B,** Subset of the metabolome data. Shown are substrate-metabolites that increase in 24 TS mutants (blue dots). Each strip of the dot plot shows one metabolite in all 94 TS mutants (mod. z-score normalized). Dots are means (n = 3). Each metabolite is the substrate for at least one TS mutant enzyme, which is indicated next to each strip of the dot plot. **C,** Pathway focused analysis of the metabolome data. Pathways with metabolite increases (mod. z-score > 3) are highlighted in the heatmap. The color code indicates if one, two, or more than two metabolites increase in a particular pathway. Target-pathways (yellow boxes) are defined as pathways that involve a TS mutant enzyme. **D,** Number of pathways that respond per strain, and number of TS mutants that led to responses in a pathway. A “response” means that at least one metabolite increases in a pathway (mod. z-score > 3). The box-whisker plot indicates the median (yellow line), and the 25^th^/75^th^ percentiles (grey box).

Overall metabolome profiles were also TS mutant-specific because there was almost no correlation between pair-wise metabolome profiles of two TS mutants (or the control strain). For example, the average Pearson correlation coefficient (PCC) across all pairs of the 95 metabolomes was 0.02 (Fig EV 3). Only 29 pairs of TS mutants had similar metabolome profiles (PCC > 0.7, Suppl Table 5), and most of these pairs were functionally related (Suppl Table 5). Ten similar pairs of TS mutants affected enzymes or regulators of the same metabolic pathway, such as PurK^I213D^ and PurE^I29W^ in the purine pathway. Two TS mutant pairs with similar metabolomes were subunits of the same enzyme complex: carbamoyl-phosphate synthetase subunits CarA^F266P^ and CarB^V322D^, and the subunits of 3-isopropylmalate dehydratase LeuC^V125P^ and LeuD^I73D^. Moreover, the TS mutant of the transcriptional regulator of the cysteine pathway CysB^V171Q^ had a similar metabolome as the TS mutants CysH^V49Q^ and CysE^V226A^, which are two enzymes in the cysteine pathway (Fig 3B). Thus, metabolome responses in the TS mutants were strong and specific, and TS mutants with similar function had similar metabolomes.

66 out of 94 TS-mutations affected enzymes that catalyze reactions in 44 metabolic pathways (Fig 3A). If these TS enzymes are less active at 42°C, they may create a metabolic bottleneck that limits flux through the associated pathway. Previous studies already showed that metabolic bottlenecks and perturbations in enzyme capacity can increase the concentration of upstream metabolites, especially the levels of substrate metabolites (Donati *et al*, 2021; Fuhrer *et al*, 2017; Fendt *et al*, 2010). Consistent with perturbations in enzyme capacity, the MurE^W381Q^ strain had a strong increase of the MurE-substrate (UDP-N-acetyl-α-D-muramoyl-L-alanyl-D-glutamate), and the HisC^I336D^ strain had an increase of the HisC-substrate (3-(imidazol-4-yl)-2-oxopropyl phosphate). Thus, the metabolome data confirmed that the proximate cause of growth defects of the MurE^W381Q^ and HisC^I336D^ strains (Fig 2C) are bottlenecks in peptidoglycan and histidine biosynthesis, respectively. In total, substrate metabolites increased in 24 strains with TS enzymes (Fig 3B). The remaining 42 strains with TS enzymes showed no increases of the direct substrates, either because substrate metabolites were not covered by the metabolome data or because the substrate metabolite is unstable, like in the case of ProA^M277P^ (Smith *et al*, 1984), PurD^V335Q^ (Cheng *et al*, 1990), and PurE^I29W^ (Mueller *et al*, 1994). Another explanation for the absence of increasing substrates is that they are used by branching or competing metabolic pathways.

Next, we examined global metabolome changes and counted the number of metabolites that increased per metabolic pathway and per TS mutant (Fig 3C, Suppl Table 6). 36 TS mutants showed a response in their “target-pathway”, which is the metabolic pathway that involves the TS enzyme. These 36 TS mutants with local responses in the target-pathway included the 24 strains with increases of direct substrates (Fig 3B) and another 12 mutants, in which other metabolites of the target-pathway increased. Apart from local responses in the target-pathways, we observed responses in distal pathways: a single TS mutation perturbed on average 12 metabolic pathways (Fig 3D), demonstrating that a single perturbation has global effects on metabolism.

Global metabolome changes can originate from regulatory interactions or from metabolites that participate in multiple pathways. For example, phosphoenolpyruvate (PEP) participates in glycolysis but also in aromatic amino acid biosynthesis. Therefore, the GapA^V17W^ strain, which has a temperature-sensitive glycolysis enzyme (glyceraldehyde-3-phosphate dehydrogenase, GapA), had a primary bottleneck in glycolysis and secondary bottleneck in aromatic amino acid biosynthesis. The glycolysis bottleneck was evidenced by increases of the substrate metabolite glyceraldehyde-phosphate (Fig EV 4), and the bottleneck in aromatic amino acid biosynthesis was evidenced by increases of shikimate-phosphate (Fig EV 4). Shikimate-phosphate and PEP are both substrates of the 6^th^ step of aromatic amino acid biosynthesis catalyzed by 3-phosphoshikimate 1-carboxyvinyltransferase (AroA), and therefore the low PEP levels in the GapA^V17W^ strain (Fig EV 4) seem to perturb the AroA capacity.

Regulatory interactions are another source of global metabolome changes in the TS mutants. For example, increases of substrate-metabolites can allosterically inhibit enzymes in another metabolic pathway. An example of this regulatory crosstalk is the PanB^L42Q^ strain, in which local metabolome changes in the phosphopantothenate pathway propagate into the tyrosine biosynthesis pathway via an allosteric interaction. PanB catalyzes the first step of phosphopantothenate biosynthesis (Jones *et al*, 1993), and the PanB-substrate is 3-methyl-2-oxobutanoate, which increased in the TS mutant PanB^L42Q^ (Fig 3B). 3-methyl-2-oxobutanoate is a known inhibitor of TyrB, which explains increases of the TryB-substrate (hydroxy-phenylpyruvate) in the PanB^L42Q^ strain (Fig EV 4).

In summary, most TS mutants showed strong and specific metabolome changes, demonstrating that TS mutants have diverse metabolic states. In 24 TS mutants the direct substrate metabolite increased, presumably, because the TS mutation reduced enzyme capacity at 42°C, which in turn leads to a bottleneck in the target-pathway. Substrate metabolites increased also in distal pathways indicating secondary bottlenecks due to metabolic-coupling (e.g. perturbation of AroA in the GapA^V17W^ strain), or regulatory interactions (e.g. perturbation of TryB in the PanB^L42Q^ strain). Next, we analyzed if increases of substrates metabolites are short-lived or if they accumulate for longer periods of time.

### Production of substrate metabolites is long-lasting and tunable by temperature

Metabolome data showed that substrate-metabolites increased in 24 TS mutants (Fig 3B). Therefore, we examined if a TS mutant produces substrate-metabolites for a longer period of time and if substrate increases are tunable by temperature. To test this, we focused on homoserine which increased in the MetA^F285W^ strain and in the ThrB^F267D^ strain (Fig 3B). MetA (homoserine *O*-succinyltransferase) catalyzes the first step in methionine biosynthesis, ThrB (homoserine kinase) catalyzes the first step in threonine biosynthesis, and homoserine is the common substrate of MetA and ThrB (Fig 4A). We cultured the MetA^F285W^ strain and the ThrB^F267D^ strain at different temperatures and measured homoserine in the whole culture broth using LC-MS/MS (Fig 4B). Our LC-MS/MS method could not separate homoserine and threonine, and therefore we measured the total pool of homoserine and threonine. In the following, we assume that homoserine is responsible for the increases of the total pool of homoserine and threonine because threonine is downstream of MetA and ThrB. At 30°C, homoserine did not accumulate in both mutants (MetA^F285W^ and ThrB^F267D^, Fig 4B), demonstrating that the two TS enzymes are functional at 30°C. This is consistent with their normal growth phenotype at 30°C, which was similar to the control strain (Fig 4B). Higher temperatures induced homoserine production: gradual temperature increases (34°C, 37°C, 39°C, and 43°C) led to gradual increases of homoserine production in the ThrB^F267D^ strain (Fig 4B). At the same time, growth of the ThrB^F267D^ strain decreased at higher temperatures. At 43°C, the ThrB^F267D^ strain did not grow and stably produced homoserine for at least 6 h. The MetA^F285W^ mutant showed a similar behavior as the ThrB^F267D^ strain, but produced less homoserine.

**Figure 4.**
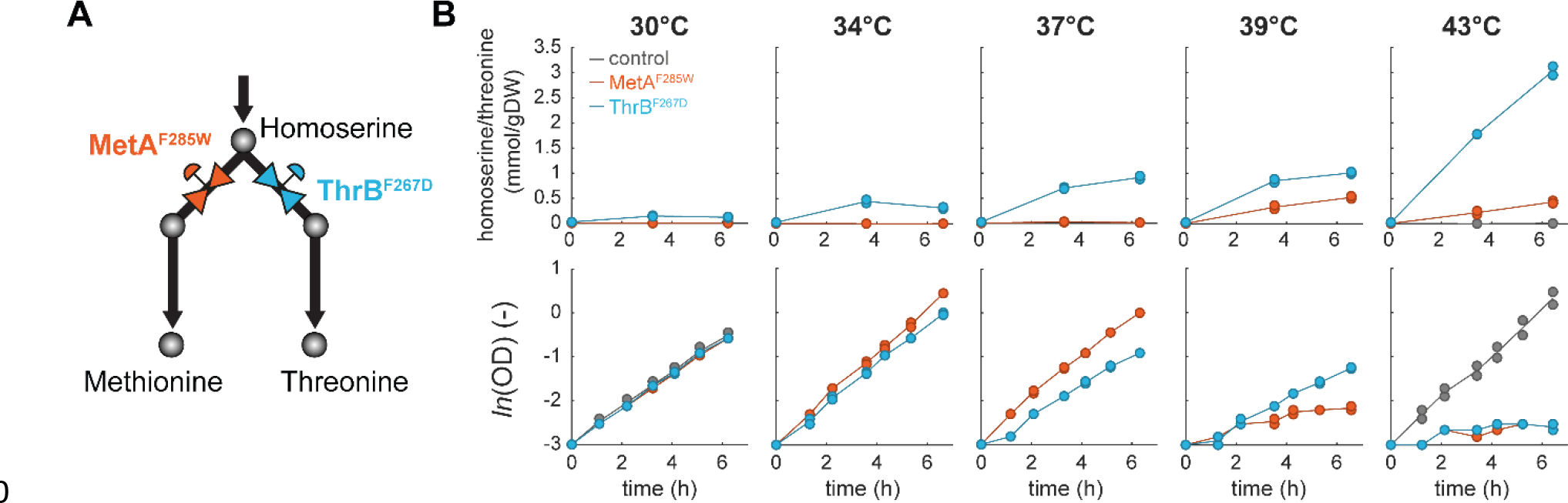
**MetA^F285W^ and ThrB^F267D^ function as metabolic valves and overproduce homoserine.** **A,** Two TS mutants, MetA^F285W^ and ThrB^F267D^, at the homoserine branchpoint. MetA catalyzes the first step in the methionine biosynthesis pathway in *E. coli*, ThrB catalyzes the first step in the threonine biosynthesis pathway. **B,** Biomass-specific concentration of the pool of homoserine and threonine (LC-MS/MS could not separate homoserine and threonine). Shown are cultures of the MetA^F285W^ strain (red), the ThrB^F267D^ strain (blue), and the control strain (grey). The strains were grown in shaking flasks at the indicated temperatures. Dots show samples from two replicates (n = 2). Lower charts show the optical density (OD) in the same cultures.

Thus, the TS mutants MetA^F285W^ and ThrB^F267D^ enable tight control of homoserine overproduction: at 30°C, they do not overproduce homoserine, whereas higher temperatures gradually increase homoserine production. Both strains remain metabolically active for at least 6 h, even if growth is fully arrested (at 43°C). In additional experiments, we found that homoserine production continued for up to 24 h, with specific production rates of 48 µmol gDW^-1^ h^-1^ for the MetA^F285W^ strain and 270 µmol g_DW_^-1^ h^-1^ for the ThrB^F267D^ strain (Fig EV 5). Combining the MetA^F285W^ and the ThrB^F267D^ mutations into a double TS mutant had an additive effect, because it increased the homoserine production even further to 477 µmol gDW^-1^ h^-1^. Other three TS mutants (LysA^L280D^, AroC^V339P^, and ArgG^L114P^) showed a similar overproduction of substrate metabolites during a phase of 24-hour growth arrest (Fig EV 5), demonstrating that TS mutants are generally applicable to produce a wide-range of bacterial metabolites.

### Temperature-sensitive DNA polymerase DnaX^L289Q^ decouples microbial growth from overproduction of arginine

Finally, we tested if we could use a TS mutant to control growth of an engineered overproduction strain. Therefore, we inserted the TS mutation DnaX^L289Q^ into an arginine overproducing *E. coli* strain (Sander *et al*, 2019) to control its growth by temperature (Fig 5A). We selected the DnaX^L289Q^ mutation, because the DnaX^L289Q^ mutant grew well between 30°C and 34°C and stopped growing at temperatures above 38°C (Fig EV 2). Moreover, the DnaX^L289Q^ mutant showed no strong metabolome changes at 42°C (Fig 3C), and we assumed that this reduces interferences with arginine production. The arginine overproduction strain was constructed by removing transcriptional feedback of the arginine repressor (Δ*argR*), which results in overexpression of arginine enzymes (Sander *et al*, 2019). Additionally, a point mutation was inserted into the first enzyme of the arginine pathway (ArgA^H15Y^) to remove allosteric feedback inhibition by arginine. To facilitate transport of arginine, the arginine exporter ArgO was overexpressed from a plasmid (Fig 5A). This resulted in a strain with 4 modifications: (1) overexpression of ArgO, (2) a point mutation ArgA^H15Y^, (3) a gene deletion Δ*argR* and (4) a point mutation DnaX^L289Q^. In the following, we refer to this strain as the TS arginine production strain.

**Figure 5.**
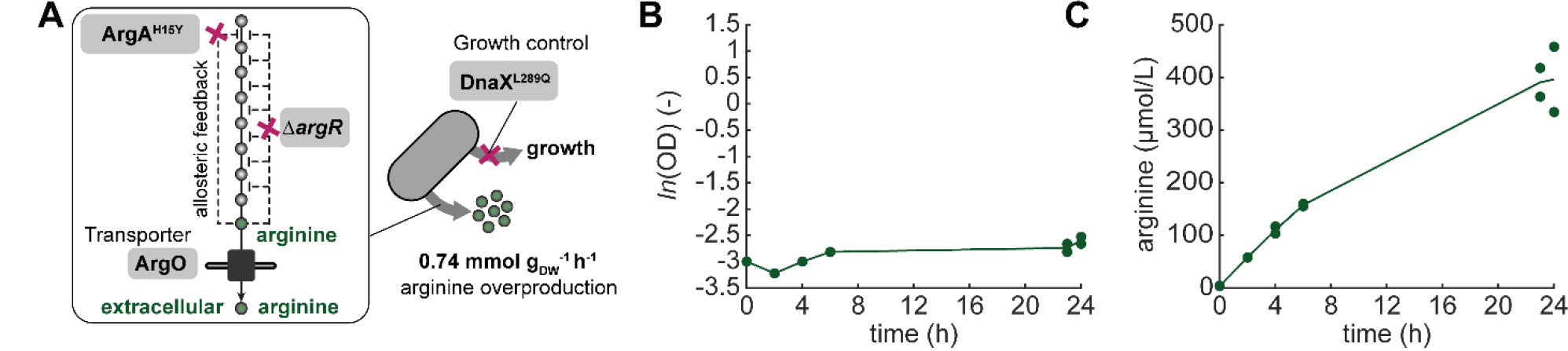
**Decoupling growth from arginine overproduction by DnaX^L289Q^.** **A,** Arginine overproduction strain with the TS mutation DnaX^L289Q^ for growth control. Dysregulation of the arginine pathway was achieved by deleting *argR* (removes transcriptional feedback) and inserting the ArgA^H15Y^ mutation (removes allosteric feedback). The arginine exporter ArgO was overexpressed from a plasmid. **B,** OD of the engineered arginine overproduction strain during cultivations at 42°C in shaking flasks. Dots are samples from two replicates (n = 2), and the line connects the means. **C,** Arginine concentration (µmol/L) during the same cultivation (shown in **B**). Arginine levels were quantified in the whole culture broth by LC-MS/MS and calibrated with an authentic arginine standard. Dots show samples from two replicates (n = 2), and the line connects the means.

The TS arginine production strain was cultivated for 24 h at 42°C in minimal glucose medium. As expected, the strain did not grow at 42°C due to the TS mutation DnaX^L289Q^ (Fig 5B). Arginine was quantified in the culture broth by LC-MS/MS to estimate the biomass specific arginine production rate (Fig 5C). The arginine production rate was 0.74 mmol gDW^-1^ h^-1^, which is 32% of the production rate (2.3 mmol gDW^-1^ h^-1^) that can be achieved with the growing arginine production strain without the DnaX^L289Q^ mutation (Sander *et al*, 2019). Thus, despite the growth arrest of the TS arginine production strain, this strain maintained a very high flux through the arginine pathway (429% of the arginine flux in growing wild-type *E. coli*), which opens up new possibilities for a temperature-controlled production of arginine in two-stage bioprocesses.

## Discussion

CRISPR-based genome editing methods have become highly efficient and versatile in their application. They enabled the construction of thousands of targeted genomic edits in large mutant libraries, which were then screened for various phenotypes such as responses to antibiotics (Dewachter *et al*, 2023; Meier *et al*, 2022; Garst *et al*, 2017) or improved production of biochemicals (Liang *et al*, 2017). Here, we used CRISPR-based genome editing to screen for single amino acid changes that cause temperature-sensitivity. So far, TS mutants were mostly constructed by random mutagenesis approaches and subsequent screening for TS growth of single colonies (Ben-Aroya *et al*, 2008; Kofoed *et al*, 2015; Li *et al*, 2011). Our approach provides a more systematic way to construct TS mutants, and should therefore facilitate the discovery of TS mutants in other organisms. We designed and introduced 15,120 mutations that resulted in up to 50 single amino acid changes in 346 essential proteins (out of all 352 essential proteins). Amino acid changes were designed with the TSpred algorithm (Tan *et al*, 2014), such that they have a high chance to decrease thermal stability of the target protein. Our data indicate that many of these predictions decreased protein stability already at 30°C, because a large fraction of the CRISPR library (55%) had strong fitness defects at 30°C. Nevertheless, 1,269 strains in the CRISPR library grew well at 30°C and showed fitness defects at 42°C, thus indicating that they are temperature-sensitive.

We reconstructed a panel of 94 TS mutants and showed that they function as growth switches, which arrest cellular growth at temperatures between 32°C and 44°C. A unique advantage is that some of these growth switches operate at very fast time scales, as exemplified for the MurE^W381Q^ and HisC^I336D^ mutants that switched from growth to no growth within 30 min. Future studies should explore if all mutants respond at such fast time scales, and if the growth switches are reversible (i.e., if cells re-grow upon temperature decreases). Others constructed growth switches by expressing essential genes under control of inducible promoters (Izard *et al*, 2015) or by repressing transcription of essential genes by CRISPR interference (Li *et al*, 2020, 2016). These growth switches operate at slower time scales than TS mutants, because they interfere with *de novo* synthesis of essential proteins and do not alter the activity of proteins that are already expressed. The consequence is that knockdowns with CRISPR interference, for instance, show significant time delays between induction and the emergence of a phenotype (Anglada-Girotto et al, 2022; Donati et al, 2021).

A further challenge for all growth switches is that bacteria can escape the growth arrest by mutations and other compensatory mechanisms. For example, cells can escape a CRISPRi-mediated growth arrest by loss-of-function mutations in the dCas9 or sgRNA sequences that alleviate the transcriptional repression. We expect that TS mutants are less prone to escaping their growth-arrested state, because this would require unique mutations that restore the protein function at high temperatures. Moreover, combining multiple TS mutations (e.g., our double TS mutant MetA^F285W^ ThrB^F267D^) should further decrease the risk of escape mutations, since cells have to restore the function of two essential proteins.

The TS mutants allowed us to arrest growth in various ways, e.g., by blocking replication (DnaX^L289Q^) or metabolic functions like histidine biosynthesis (HisC^I336D^). The way we arrested growth had strong effects on metabolism under growth arrest: some TS mutants had a wild-type-like metabolome, whereas other TS mutants had strong metabolome changes in many metabolic pathways. This gene-specific response of metabolism of the TS mutants might be relevant for their application in bioprocesses, where TS mutants could function as metabolic valves (Venayak *et al*, 2018) or to control growth and production phases in two-stage bioprocesses (Burg *et al*,

2016). So far, it remains unclear if some TS mutants are better suited for bioprocesses than others. Maybe the choice of the TS mutant has no significant impact on titers, rates, and productivity of a process, but it seems likely that the metabolic state of a TS mutant affects these parameters. Therefore, more work is required to understand which TS mutants can improve the production of a particular bio-based product (Jang *et al*, 2023).

In conclusion, there is a growing interest in non-growing bacteria, and we have shown that TS mutants are a robust and versatile tool to control bacterial growth. Our data indicate that the metabolic state of non-growing bacteria is diverse, which means that non-growing bacteria do not enter a universal stand-by mode. Instead, the metabolic state of non-growing bacteria depends on the cellular process that causes growth arrest, and our ability to control these processes by exogenous signals like temperature will open up novel applications in metabolic engineering and industrial biotechnology.

## Methods

### M.1 Construction of plasmids

The CRISPR-Cas9 genome editing method was a modified version of the CREATE method (Garst *et al*, 2017). Two plasmids (pTS040 and pTS041) were constructed using Gibson assembly. pTS040 had the p15A origin of replication and carried a chloramphenicol resistance gene, a cassette with the homology arm for recombination, and the guide RNA of the CRISPR system under control of a constitutive promoter (PJ23119). pTS041 had the pSC101 origin of replication and carried a kanamycin resistance gene, a gene for the anhydrotetracycline (aTc)-sensitive repressor *tetR*, *cas9* under control of the aTc controlled PLtetO1 promoter, the arabinose-sensitive repressor *araC*, and the *Escherichia* virus lambda genes *red* under control of the arabinose-controlled promoter ParaBAD. pTS055 was pTS040 with a spectinomycin resistance gene instead of a chloramphenicol resistance gene. pTS056 had a p15A origin of replication, an ampicillin resistance gene, the anhydrotetracycline(aTc)-sensitive repressor *tetR*, and *argO* encoding for an arginine exporter under the aTc controlled PLtetO1 promoter. pTS056 was based on a plasmid from (Sander *et al*, 2019). Plasmids were constructed with Q5 High-fidelity DNA polymerase (New England BioLabs Inc., NEB) and the Gibson Assembly Master Mix (NEB). We used the DNA Clean & Concentrator Kit (Zymo Research) to purify DNA after PCRs.

### M.2 Design of the temperature sensitive E. coli library

The TSpred tool (Tan *et al*, 2014; Varadarajan *et al*, 1996) was used to predict TS mutations for all 352 genes that are essential for growth of *E. coli* on minimal glucose medium (Goodall *et al*, 2018; Patrick *et al*, 2007). It is predicted, which amino acid of a protein, upon substitution by one of the five amino acids alanine, tryptophan, glutamine, aspartate, or proline, is likely to introduce temperature-sensitivity. If possible, crystal structures of the target proteins were given as an input for the algorithm. Otherwise, amino acid sequences were given as input. The predictions are listed in Suppl Tab 1.

Because the quality of a protospacer for CRISPR-Cas9 genome editing is strongly affected by the distance of the PAM to the target site (Garst *et al*, 2017) and its off-targets, we considered every PAM within 30 bp distance for every predicted site and tested if a silent PAM-mutation was possible. In some cases, the PAM and the target site overlapped, and, therefore, not every amino acid substitution was possible to remove the PAM. The Cas-OFFinder (Bae *et al*, 2014) was used to identify off-targets with up to 4 mismatches. We also tested if the protospacers have a 11 PAM-proximal perfect match to multiple PAMs (Rousset *et al*, 2018). Based on these results, we then ranked each available design for every site with a custom scoring system and chose 10 predicted sites for each gene that had the highest-ranking designs (Methods Table 1). We excluded designs that did not reach a minimum score of 3 such that some target genes yielded no or less than 10 designs. The final library contained 16,038 members covering 346 genes.

**Methods Table 1.** **Custom scoring system of cassette designs.** The distance between the silent PAM-mutation and the target site is considered in steps of 3 nucleotides (nt). Cassette designs in “class 1” do not have another 11 nt PAM-proximal perfect match and no off-target with up to 4 mismatches. Cassette designs in “class 2” do not have another 11 nt PAM-proximal perfect match. Cassette designs in “class 3” do not have an off-target with up to 4 mismatches. Cassette designs in “class 4” have other 11 nt PAM-proximal perfect matches and off-targets with up to 4 mismatches. For each site, the ten highest ranking cassette designs were chosen. The minimum score was 3. If only 4 amino acid substitutions were possible at a given site, a penalty of -0.95 was applied to the score (3 substitutions: -1.25, 2 substitutions: -3).

| distance | 0 nt | 3 nt | 6 nt | 9 nt | 12 nt | 15 nt | 18 nt | 21 nt | 24 nt | 27 nt | 30 nt |
| --- | --- | --- | --- | --- | --- | --- | --- | --- | --- | --- | --- |
| class 1 | 5 | 4.8 | 4.6 | 4.4 | 4.2 | 4.0 | 3.8 | 3.6 | 3.4 | 3.2 | 3.0 |
| class 2 | 4.41 | 4.21 | 4.01 | 3.81 | 3.61 | 3.41 | 3.21 | 3.01 | 2.81 | 2.61 | 2.41 |
| class 3 | 3.9 | 3.7 | 3.5 | 3.3 | 3.1 | 2.9 | 2.7 | 2.5 | 2.3 | 2.1 | 1.9 |
| class 4 | 2 | 1.8 | 1.6 | 1.4 | 1.2 | 1.0 | 0.8 | 0.6 | 0.4 | 0.2 | 0 |

Oligonucleotides to construct the library were 200 bp long and contained in the following order: a spacer sequence (‘TCCTCTGGCGGAAAGCC’), a homology sequence with the desired mutation and a silent PAM-mutation, another spacer (‘GATC’), the J23119 promoter (‘TTGACAGCTAGCTCAGTCCTAGGTATAATACTAGT’), a protospacer, and a part of the sgRNA-Cas9 handle (‘GTTTTAGAGCTAGAAATAGCAAGTTAAAATAAGGCTAG’).

### M.3 Strain construction

#### M.3.1 Cloning the CRISPR libraries

The oligonucleotide pools were manufactured by Twist Bioscience (South San Francisco, United States). The oligonucleotides were used as template for PCR amplification (oligonucleotide concentration: 0.1 µM; 15 cycles). The PCR products of correct size were purified by agarose gel electrophoresis (NucleoSpin Gel and PCR Clean-up Kit, Macherey-Nagel). The purified linear DNA was used for cloning of pTS040 by Gibson assembly (NEBuilder HiFi DNA Assembly Reaction, NEB) and electroporation of *E. coli* MegaX DH10B T1^R^ cells (Invitrogen, Thermo Fisher Scientific Inc.).

*E. coli* BW25113 carrying pTS041 was cultured in LB medium at 37°C under shaking of 220 rpm until exponential growth. Expression of the *Lambda red* genes was induced with L-arabinose (7.5 g/L). After 30 min, the culture was harvested for electroporation with the pooled pTS041 plasmid library (0.1 cm Gene Pulser Cuvette #165-2089 and Micropulser, BioRad). Cells were recovered in SOC medium with kanamycin and 1 µM aTc for Cas9 induction at 30°C for 2 h and streaked out on to LB agar plates with kanamycin, chloramphenicol, and 1 µM aTc. After incubation overnight at 30°C, colonies were pooled by flushing the agar plates with LB medium, glycerol added (final concentration: 22 vol.-%), the OD was determined, and the strain library stored as cryo stocks.

### M.3.2 Construction of the double TS mutant MetA^F285W^+ThrB^F267D^ by sequential CRISPR-Cas9 genome editing

A 50 mL LB culture without chloramphenicol of the TS mutant MetA^F285W^ (*E. coli* BW25113 *metA*(F285W)// pTS041// pTS040(MetAF285W)) was started from cryo stock and incubated for 22 h at 30°C under 220 rpm of shaking. After diluting the culture 1:10,000 in fresh LB without chloramphenicol, the cells were further incubated overnight at 30°C under shaking of 22 rpm. A fresh culture was started in the morning by 1:50 dilution in the same medium and conditions. After ca. 2 h, *Lambda Red* was induced for 30 min by addition of L-arabinose (7.5 g/L). Cells were subsequently transformed by electroporation with pTS055(ThrB^F267D^) (0.1 cm Gene Pulser Cuvette #165-2089 and Micropulser, BioRad). Cells were recovered in SOC medium with kanamycin and 1 µM aTc for Cas9 induction at 30°C for 2 h and streaked out on to LB agar plates with kanamycin, spectinomycin, and 1 µM aTc. After incubation at room temperature, single isolates were stored as cryo stocks and checked for correct mutations by Sanger sequencing.

#### M.3.3 Construction of the DnaX^L289Q^ arginine overproduction strain

The strain *E. coli* MG1655 Δ*argR* ArgA-H15Y//pTS041 was used for transformation with pTS040(DnaX^L289Q^) as described in M.3.1 and M.3.2. The genomic edit was confirmed by Sanger sequencing. Subsequently, the strain was transformed with pTS056 that was used to overexpress the arginine exporter ArgO. It was previously described that basal expression of ArgO was sufficient (Sander *et al*, 2019) such that aTc was not added to subsequent cultures.

### M.4 Cultivations

If not stated otherwise, minimal medium (M9) was used for the experiments and contained 42.2 mM Na2HPO4, 11.3 mM (NH4)2SO4, 22 mM KH2PO4, 8.56 mM NaCl, 1 mM MgSO4 x 7 H2O, 100 µM CaCl2 x 2 H2O, 60 µM FeCl3, 6.3 µM ZnSO4 x 7 H2O, 7 µM CuCl2 x 2 H2O, 7.1 µM, MnSO4 x 2 H2O, 7.6 µM CoCl2 x 6 H2O, and 2.8 µM thiamine-HCL. 5 g/L glucose was used as carbon source. M9 and LB agar plates contained 1.5% agar. 30 µg/L chloramphenicol, 50 µg/mL kanamycin, 100 µg/mL carbenicillin, and 50 µg/mL spectinomycin were added to the media when required.

#### M.4.1 Competition experiment and sampling for amplicon sequencing

The TS plasmid library (before electroporation of *E. coli* BW25113//pTS041) was used as a sample for amplicon sequencing (“sample before recombination”). Plasmids were extracted from the cryo stock of the TS strain library (*E. coli* BW25113//pTS041//pTS040(TS-library), “sample after recombination”). 75 mL M9 medium was inoculated with 200 µL of the TS strain library from cryo stock and incubated in a 500 mL shake flask for 15 h at 30°C under shaking of 220 rpm. 10 mL of the exponentially growing culture were used for plasmid extraction (“sample time point zero”). 300 mL of M9 medium was inoculated with the previous culture to a start OD of 0.1. The 300 mL culture was split up to each 150 mL for cultivation in 1 L-shake flasks at 30°C and 42°C under shaking of 220 rpm. Every 3 h, the 150 mL cultures were back diluted to an OD of 0.1. Every 2 h, a sample for plasmid extraction was taken (sample volume x OD ≥ 5).

#### M.4.2 Plate reader cultivations

500 µL of LB in 2 mL deep well plates (96-well) were inoculated from cryo stocks, covered with Breathe-Easy (Diversified Biotech BEM-1) adhesive membrane, and incubated for 6 h at 30°C under shaking at 220 rpm. 500 µL M9 (5 g/L glucose) were inoculated with 1 µL of the LB precultures and incubated overnight at 30°C in 2 mL deep-well plates under shaking of 220 rpm. 297 µL of M9 (5 g/L glucose) were inoculated with 3 µL of the overnight cultures in 96-well Greiner plates (flat-bottomed). 150 µL were transferred to a second 96-well Greiner plate. Each plate was incubated at each two different temperatures. Epoch 2 (BioTek, now: Agilent Technologies) or Infinite 200 Pro (TECAN trading AG) plate readers were used for incubation and measurements of OD at 600 nm every 10 min. Maximum specific growth rates were calculated in exponential growth phases if applicable.

#### M.4.3 Dynamic switch from 30°C to 42°C with the TS mutants MurE^W381Q^ and HisC^I74Q^

5 mL LB cultures were started from cryo stock. After ca. 6 h at 30°C under shaking at 220 rpm, 5 mL M9 (5 g/L glucose) overnight cultures (30°C, 220 rpm) were started using 25 µL of the LB culture for inoculation. Overnight M9 cultures were washed: the cultures were pelletized by 5 min of centrifugation at 4,000 rpm and 30°C. After removing the supernatant, 5 mL of fresh M9 glucose medium was added for resuspending cells. This step was repeated further 2 times. Final 15 mL cultures were started in 100 mL shaking flasks at an OD of 0.05 and incubated for 5 h under shaking of 220 rpm and 30°C. Then, cultures were transferred to 42°C for further incubation. The OD600 was measured regularly.

#### M.4.4 96-well cultivation and sampling for metabolomics by flow-injection mass spectrometry

500 µL of LB in 2 mL deep well plates (96-well) were inoculated from cryo stocks, covered with Breathe-Easy (Diversified Biotech BEM-1) adhesive membrane, and incubated for 6 h at 30°C under shaking at 220 rpm. 495 µL M9 (5 g/L glucose) were inoculated with 5 µL of the LB precultures and incubated for 24 h at 30°C in 2 mL deep-well plates, under shaking of 220 rpm. 100 µL M9 precultures were transferred to 900 µL of fresh M9 (5 g/L glucose) in 2 mL deep-well plates and incubated for 16 h at 42°C, under shaking of 220 rpm. 850 µL of the liquid culture was centrifuged in 2 mL deep-well plates for 15 min at 4,000 rpm at 4°C. The supernatant was removed and the cell pellets stored at -80°C. 100 µL of -20°C cold 40:40:20 acetonitrile:methanol:water was added to the frozen cell pellets and incubated for 4 h at -20°C. The plate was vortexed and 80 µL of the cell extract transferred to v-bottomed 96-well storage plates. The cell extracts were stored at -80°C until further analysis by FI-MS.

#### M.4.5 Metabolic valve experiments with MetA^F285W^ and ThrB^F267D^

5 mL LB cultures were started from cryo stock. After ca. 6 h at 30°C under shaking at 220 rpm, 40 mL M9 (5 g/L glucose) overnight cultures (30°C, 220 rpm) were started using 200 µL of the LB culture for inoculation. Overnight M9 cultures were washed: the cultures were pelletized by 5 min centrifugation at 4,000 rpm and 40°C. After removing the supernatant, 30 mL of fresh M9 glucose medium was added for resuspending cells. This step was repeated further 2 times. Final 80 mL cultures were started at an OD of 0.05, and split-up to five 15 mL cultures in 100 mL shaking flasks. Each of the five 15 mL cultures were incubated at a different temperature (30/34/37/39/43°C) for ca. 6 h under shaking of 220 rpm. At the start of the cultivation and hourly throughout the cultivation, the OD600 was measured. At the start of the cultivation and after 3 h and 6 h, whole culture broth samples for LC-MS/MS analysis were taken.

#### M.4.6 Two-stage production experiments (24 h)

5 mL LB cultures were started from cryo stock. After ca. 6 h at 30°C under shaking at 220 rpm, 5 mL M9 (5 g/L glucose) overnight cultures (30°C, 220 rpm) were started using 25 µL of the LB culture for inoculation. Overnight M9 cultures were washed: the cultures were pelletized by 5 min of centrifugation at 4,000 rpm and 40°C. After removing the supernatant, 5 mL of fresh M9 glucose medium was added for resuspending cells. This step was repeated further 2 times. Final 15 mL cultures were started in 100 mL shaking flasks at an OD of 0.05 and incubated for 24 h under shaking of 220 rpm and 42°C. At the start of the cultivation and after 2 h, 4 h, 6 h, and 24 h incubation, the OD600 was measured, and whole culture broth samples for LC-MS/MS analysis were taken.

#### M.4.7 Arginine overproduction experiment with a DnaX^L289Q^ mutant

10 mL LB cultures were started from cryo stock. After ca. 6 h at 30°C under shaking at 220 rpm, 10 mL M9 (5 g/L glucose) overnight cultures (30°C, 220 rpm) were started using 50 µL of the LB culture for inoculation. Overnight M9 cultures were washed: the cultures were pelletized by 5 min centrifugation at 4,000 rpm and 40°C. After removing the supernatant, 10 mL of fresh M9 glucose medium was added for resuspending cells. This step was repeated an additional 2 times. The final 50 mL cultures were started in 500 mL shaking flasks at an OD of 0.05 and incubated for 24 h under shaking of 220 rpm and 42°C. At the start of the cultivation and after 2 h, 4 h, 6 h, 23 h, and 24 h incubation, the OD600 was measured, and whole culture broth samples for LC-MS/MS analysis were taken.

### M.5 Sample processing for NGS

Using 3 ng total plasmid DNA, a plasmid part covering the homology sequences and protospacers was amplified (15 cycles) using two primers suited for further indexing PCRs (forward: ‘TCGTCGGCAGCGTCAGATGTGTATAAGAGACAGGTATCACGAGGCAGATCCTCTG’,reverse: ‘GTCTCGTGGGCTCGGAGATGTGTATAAGAGACAGACTCGGTGCCACTTTTTCAAGTT’). Amplicons were purified by AMPure XP PCR beads (Beckman Coulter, #A63881). Using standard Illumina indexing primers, amplicons were indexed in a second PCR and again purified by bead-clean up. Amplicons were pooled and sequenced on an Illumina NextSeq500 (paired-end, NextSeq™ 500 Mid Output Kit v2.5, #20024908, 300 cycles). Two cartridges were required to yield the desired sequencing depth of around 4 million reads per sample.

### M.6 NGS data analysis

Demultiplexed paired-end reads were aligned, merged (based on overlapping sequences), and trimmed to the region of interest using a custom Matlab script. The resulting processed reads were mapped against the designed sequences of the library. For each library member, the number of matching reads was counted. Only reads that shared a 100% identity with a designed sequence were counted since mutations could indicate a malfunction of the CRISPR-Cas9 genome editing system with no genomic edit. Fitness scores were calculated by normalizing the read count of an individual mutant to the total number of reads of a sample and by subsequently normalizing the data to the first sample of the experiment (t = 0 h). Using the fitness scores, the area under the curve (*AUC*) was determined for the 30°C and 42°C time series of each mutant (*i*). An error *e* was estimated using the fitness scores *n̅* for each replicate (*A* and *B*) and time point *t* normalized to the mean fitness scores:

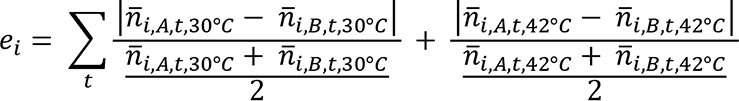

### M.7 Scoring temperature-sensitivity

Only mutants were considered that met following criteria:

– They had at least an average of 15 reads at time point zero (*r̅*_*t* =0*h*_).
– The mean fitness score of the last sample at 30°C (*n̅*_*i*,*t*=12ℎ,30°*C*_) was greater than 0.3.
– The mean fitness score of the last sample at 42°C (*n̅*_*i*,*n*=12ℎ,42°*C*_) was lower than 0.4.
– The error *e*_*i*_ was lower than 15.
– The area under the curve for the 30°C time series (*AUC*_*i*,30°*C*_) was greater than 5.
– The mean fitness score of the last samples for the different temperatures fulfilled following criterion:

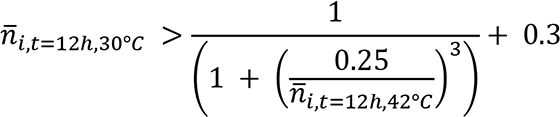

For each gene, the putatively temperature-sensitive mutants were sorted by the number of reads at time point zero *r̅*_*t=0ℎ*_, the relative area under the curve difference 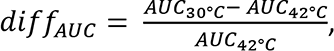 the difference between the fitness scores at 30°C and 42°C at the last time point (*diff*_*t*=12_), and the error *e*. Based on placement in the sortings (*rank*), a *score* was calculated for each candidate

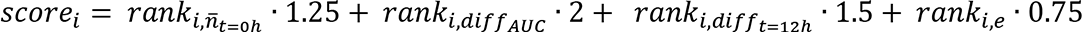

For each gene, the putatively temperature-sensitive mutant with the lowest *score* was selected for a new library with a total of 250 mutants.

### M.8 Metabolomics

#### M.8.1 Flow-injection mass spectrometry (FI-MS)

Flow-injection mass spectrometry (FI-MS) was performed as described before (Fuhrer *et al*, 2011; Farke *et al*, 2023). An Agilent 6546 QTOF mass spectrometer (Agilent Technologies, Santa Clara, USA) was used to analyze metabolite levels in metabolite extracts. The source parameters were: source gas 225°C, flow rate of the drying gas 11 L/min, nebulizer pressure 20 psi, sheath gas temperature 350°C, sheath gas flow 10 L/min, nozzle voltage 2000 V. Spectra in a 50-1100 m/z range were acquired in 10 Ghz mode with an acquisition rate of 1.4 spectra/s. The mobile phase was 10 mM (NH4)2CO3, 0.04 % NH4OH, 60:40 Isopropanol:H2O. The reference masses for online mass calibration in negative mode were 59.050 Da (C3H8O, Isopropanol) and 1033.988 (C18H18F24N3O6P3, HP-921); in positive mode, 121.050 Da (C5H4N4, Purine) and 922.009 Da (C18H18F24N3O6P3, HP-921).

#### M.8.2 FI-MS data analysis

Raw data files were converted into “.mzXML” files by MSConvert (Chambers *et al*, 2012). Following data analysis was performed by custom MATLAB scripts that utilized MATLAB functions (The MathWorks, Inc., Massachusetts, USA). The 10 spectra with the highest signal in the total ion count (TIC) were summed. Peaks with a minimum peak height of 1000 units and a peak prominence of 500 units were selected, and annotated with a 3 mDa tolerance by matching monoisotopic masses of metabolites with a single proton loss for negative mode and single proton gain in positive mode. Double annotations (positive and negative mode) were manually cured based on peak shape and height. For each metabolite, the maximum height of the annotated peak was taken for further analysis. The data was then normalized to the control strain and converted into log2 space. Subsequently, modified z-scores for each metabolite were calculated as following:

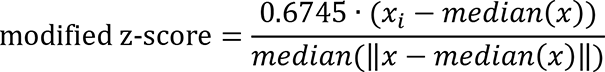

#### M.8.3 Targeted metabolomics by LC-MS/MS

Whole culture broth samples were taken by transferring 100 µL of the culture broth to -20°C cold 50:50 acetonitrile:methanol in 1.5 mL reaction tubes. The samples were stored at -80°C until further processing. The samples were centrifugated for 15 min at 17,000 g and -9°C. Metabolite concentrations in the supernatant were analyzed by an isotope-ratio based LC-MS/MS method (Guder *et al*, 2017). Changes to the LC parameters were: in the initial 0.3 min, the analyte was discarded into the waste. Between 0.3 and 2.0 min the analyte was injected to the ESI. 2.0 to 2.3 min the analyte was discarded. An internal, fully ^13^C-labelled standard was calibrated with authentic ^12^C-metabolite standards. Based on the calibrated ^13^C-standard and isotope ratios, absolute metabolite concentrations in the samples were calculated. Homoserine and threonine could not be distinguished. We used an authentic homoserine standard to calculate absolute concentrations.

## Acknowledgments

This work was supported by the ERC starting grant 715650. Amplicon next generation sequencing was performed and supported by Janina Geißert and the NGS Competence Center Tübingen (NCCT) and its technology platforms. We thank Niklas Farke for discussions.

## CRediT authorship contribution statement

*Thorben Schramm:* Conceptualization, Investigation, Visualization, Project administration, Writing, Formal analysis.

*Vanessa Pahl:* Investigation (Arginine overproduction)

*Hannes Link:* Conceptualization, Supervision, Project administration, Visualization, Writing, Funding acquisition.

