## Supplemental Figures for "Mapping temperature-sensitive mutations at a genome-scale to engineer growth-switches in *E. coli*"

### Extended View Figures

### and

#### Expanded View

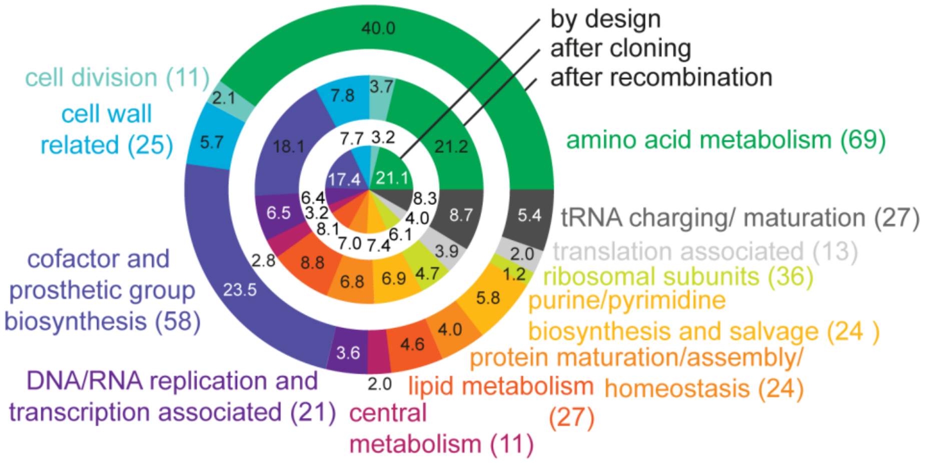


**Figure EV 1 – The CRISPR library composition at different steps of construction.**

The CRISPR library covered 346 essential genes in 12 functional categories. The chart shows the relative share of the 12 categories in the library at different steps in the construction. The inner circle indicates the composition in the original design, the middle circle the composition after cloning of the pooled plasmid library, and the outer circle the composition after inserting the mutations to the genome (recombination step).

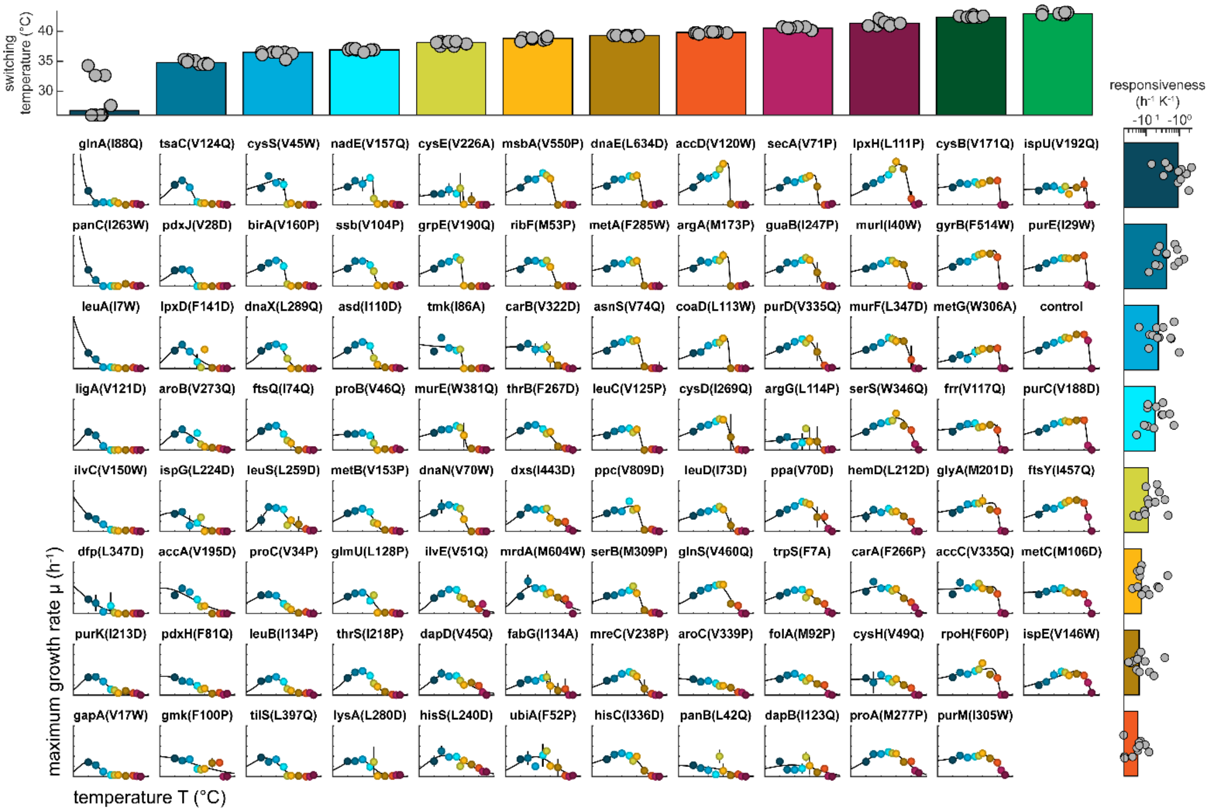


**Figure EV 2 – The maximum specific growth rates of 94 TS mutants at different temperatures.**

The charts show the maximum specific growth rates μ ($h^{-1}$) of 94 TS mutants (and a control strain) at ten different temperatures ranging from 30°C to 44°C. The growth rates were determined from growth curves in 96-well microtiter plate cultivations. Dots show the mean from n=3 replicates, black vertical lines show the standard deviation. An empirical Arrhenius-type function was fitted to the data (black lines, also see **Suppl Fig 4**). The strains were sorted according to their responsiveness and switching temperature, which are parameters based on the Arrhenius-type functions. The upper dot plot shows the switching temperatures (°C) of the strains in the columns below. The dot plot on the right side shows the responsiveness ($h^{-1}$ $K^{-1}$) of the strains in the rows. The bars indicate the medians of the responsiveness values/switching temperatures.

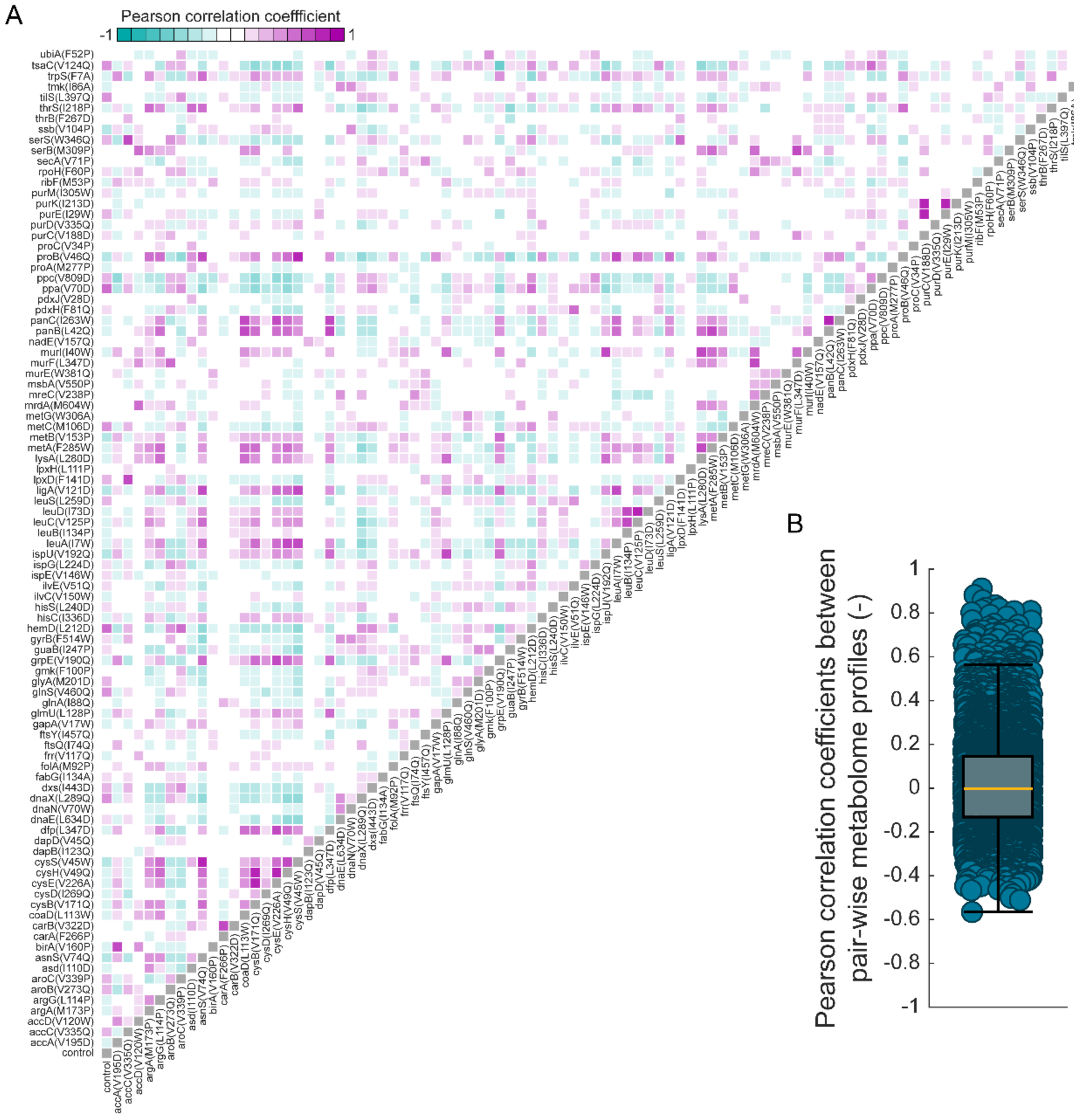


**Figure EV 3 – Correlation analysis of metabolomes of 94 TS mutants.**

**A,** The heatmap shows the Pearson correlation coefficients (PCC) between metabolite data of all pair-wise combinations of 94 TS mutants and a control strain. The metabolite levels were measured by FI-MS after cultivation of the strains in 96-well microtiter plates at 42°C for 16 h (n = 3). Data of 325 metabolites (mod. z-scores) were used to calculate the PCC values.

**B,** The dot plot shows all PCC values from **A**. The box-whiskers plot indicates the median (yellow line), and the 25$^{th}$ and 75$^{th}$ percentiles.

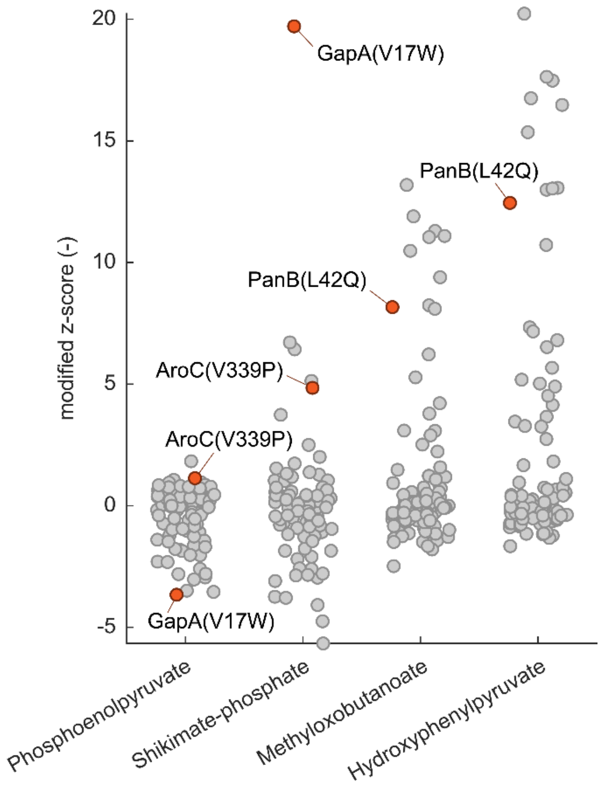


**Figure EV 4 – Metabolite increases in the TS mutants GapA$^{V17W}$, AroC$^{V339P}$, and PanB$^{L42Q}$**

The chart shows the metabolite level (mod. z-score) in the 94 TS mutants and a control strain. Grey dots are the mean (n = 3), and data of the three TS mutants GapA$^{V17W}$, AroC$^{V339P}$, and PanB$^{L42Q}$ is highlighted in red.

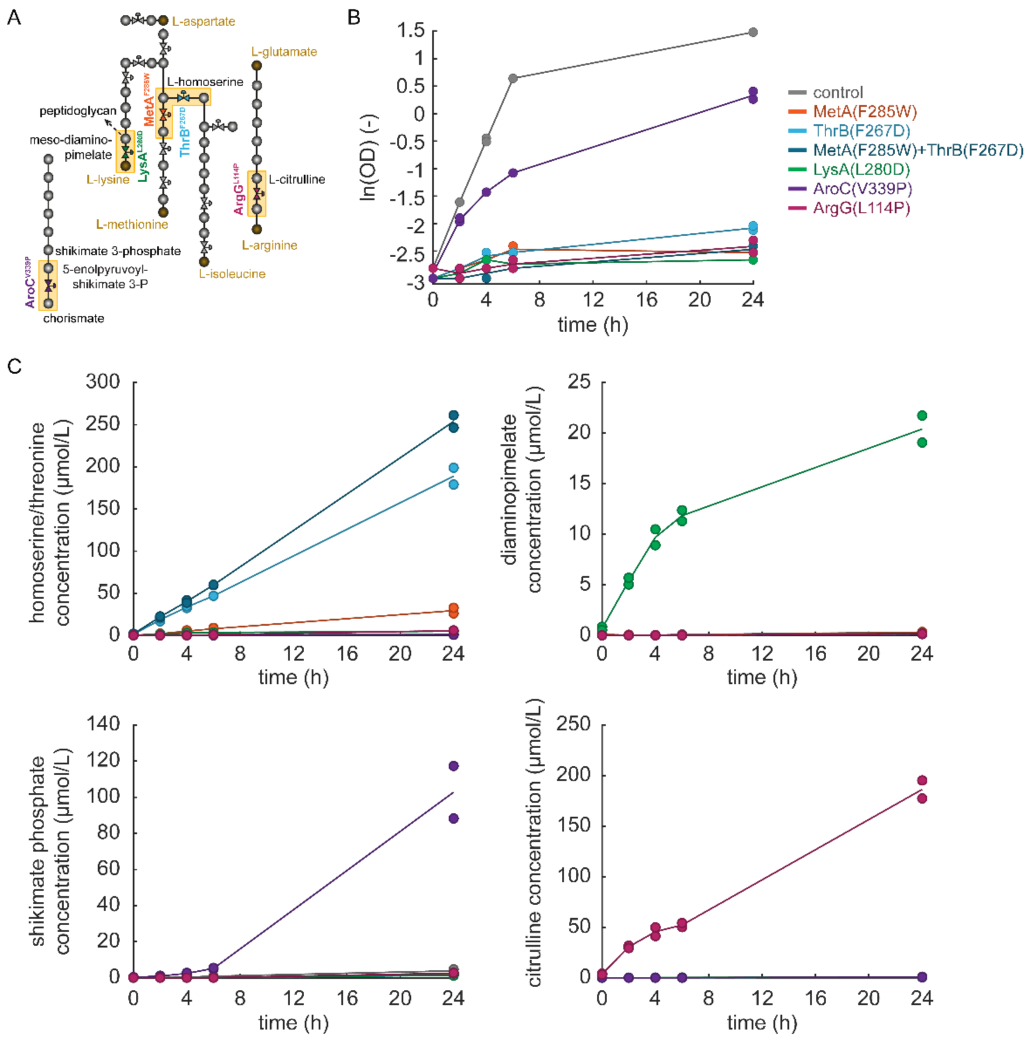


**Figure EV 5 – Substrate production in TS mutants of enzymes.**

**A,** Schematic of biosynthesis pathways of chorismate, lysine, methionine, isoleucine, and arginine. Dots represent metabolites. Valve symbols indicate TS mutant enzymes.

**B,** The chart shows the natural logarithm of biomass data (OD) from shaking flask cultivations of TS mutants and a control strain at 42°C. Dots are data from n=2 replicates, the lines connect the means.

**C,** The charts show the concentrations (µmol/L) of indicated metabolites during the cultivations from **B**. The concentrations were quantified in samples of the whole culture broth by LC-MS/MS. Dots are data from n=2 replicates, the lines connect the means.

#### Supplementary Figures

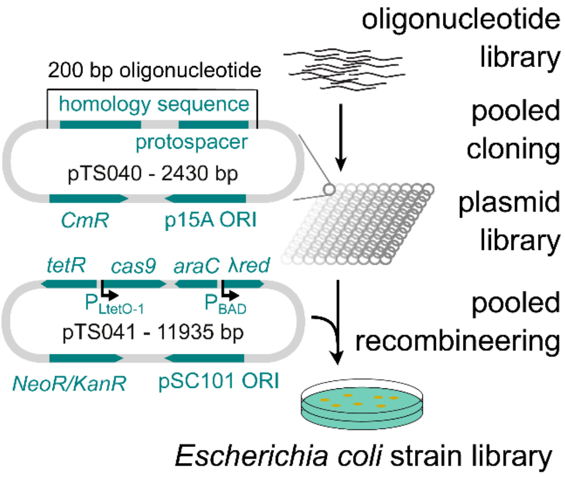


**Supplementary Figure 1 – Construction steps of pooled CRISPR strain libraries.**

Array-synthesized oligonucleotide pools were used to clone pooled plasmid libraries based on the pTS040 plasmid. pTS040 had a p15A origin of replication (ORI) and carried a chloramphenicol resistance gene as well as the homology and guide RNA sequences. In a second step, the pooled plasmid libraries were used for transformation of an *Escherichia coli* strain (BW25113) that already carried the pTS041 plasmid. pTS041 had a pSC101 ORI and carried genes for a kanamycin resistance, the transcriptional repressors *araC* and *tetR*, the *Escherichia virus Lambda red* system, and *Streptococcus pyogenes cas9*. *cas9* was under control of the $P_{LtetO-1}$ promoter. The *Lambda red* system was under control of the $P_{BAD}$ promoter. After transformation, cells were plated to agar plates. Colonies were collected from the plates and pooled yielding the final CRISPR libraries.

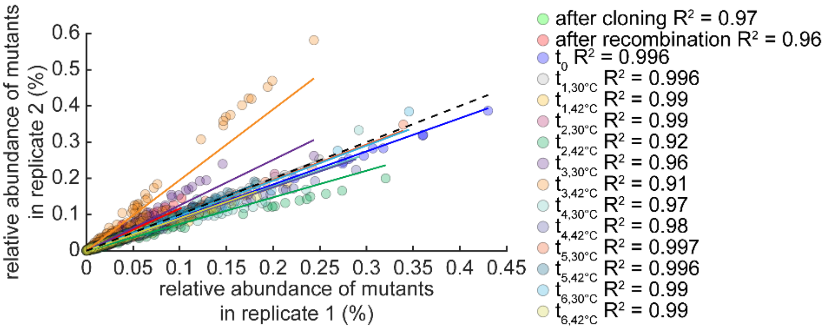


**Supplementary Figure 2 – Deep sequencing of CRISPR barcodes is reproducible.**

The parity plot shows the relative abundance of single strains in the CRISPR library at different steps during construction and during the competitive growth experiment (also see Fig 1). The abundance was measured by next generation amplicon deep sequencing (n = 2). Read counts of single mutants with 100% sequence identity were normalized to the total number of reads to calculate the relative abundance. Lines indicate linear regressions. $R^2$ is the coefficient of determination.

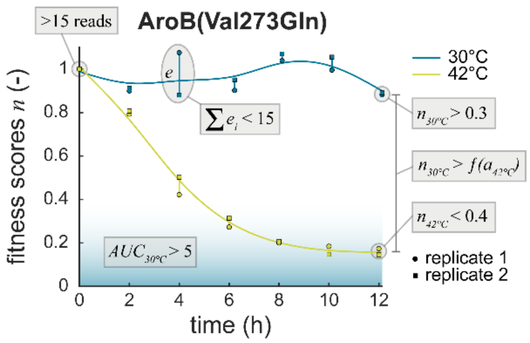


**Supplementary Figure 3 – A scoring system for TS mutations.**

Due to complex dynamics of fitness scores but also to reduce the number of putatively TS mutants, we scored temperature-sensitivity of all mutants in the CRISPR library. The chart illustrates six measures important for the scoring (also see M.7). As an example, the fitness scores of the AroB$^{Val273Gln}$ during the competitive growth experiment (Fig 1A) are shown. Squares and dots indicate replicates (n = 2). Lines are the moving average through means. Fitness scores were calculated as described in M.6. During scoring, mutants were discarded when they failed a set of minimum requirements, as indicated in the figure.

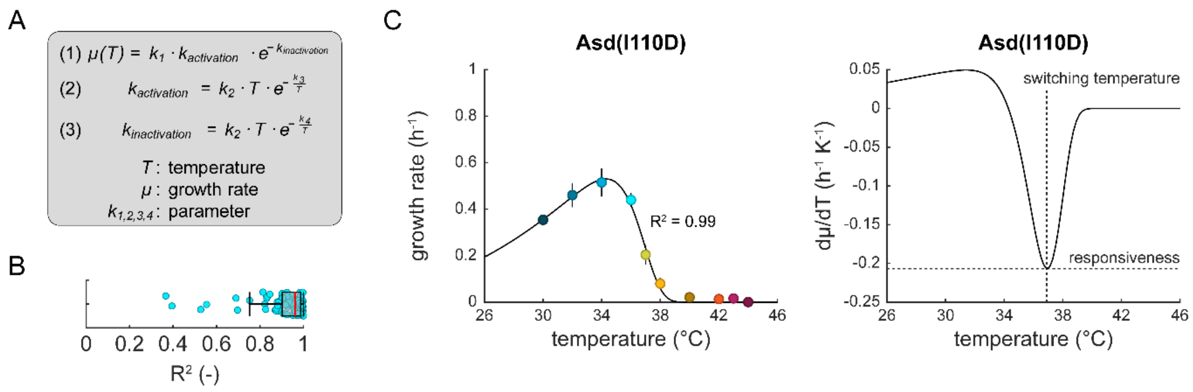


**Supplementary Figure 4 – An empirical Arrhenius-type function describes the growth rate/temperature relationship of TS mutants.**

**A,** The empirical Arrhenius-type function (1) that was used to describe the maximum specific growth rate μ of *E. coli* strains as a function of the temperature T had an activating term $k_{activation}$ (2) and an inactivating term $k_{inactivation}$ (3). The Arrhenius-type function further had four parameters $k_{1,2,3,4}$ that were fitted to experimental data.

**B,** The dot plot shows the coefficient of determination $R^2$ of the Arrhenius-type function in **A** fitted to experimental data of 94 TS mutants and a control strain without mutation. Dots are data of individual strains. The box-whiskers plot indicates the median (red line) and the 25$^{th}$ and 75$^{th}$ percentiles.

**C,** The left chart shows the maximum growth rate ($h^{-1}$) of the TS mutant $Asd^{I110D}$ at different temperatures (°C). The data was measured in microtiter plate cultivations (data also shown in Fig EV 2). Dots are the mean, vertical black lines the standard deviation (n = 3). The black line is the fitted Arrhenius-type function of **A**. $R^2$ is the coefficient of determination. The right chart shows the first derivative (dμ/dT) of the Arrhenius-type function (fitted to $Asd^{I110D}$). The minimum dμ/dT is the “responsiveness” value that also determines the “switching temperature”.

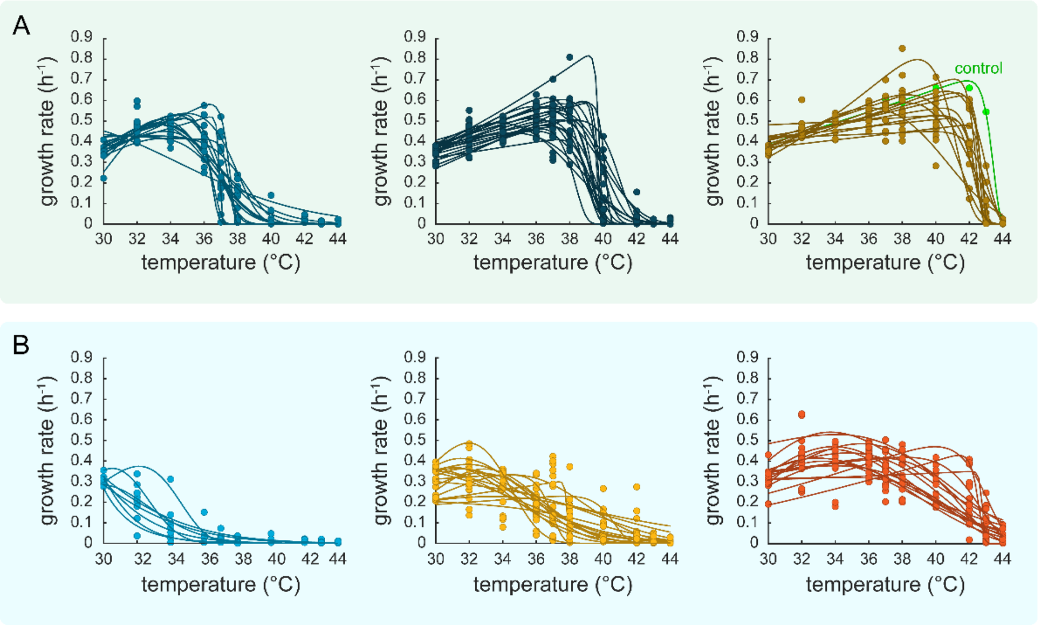


**Supplementary Figure 5 – Categorization of TS mutants based on their growth rate/temperature dependencies.**

The charts in **A** and **B** show the maximum specific growth rates ($h^{-1}$) of 94 TS mutants and a control strain without mutation at ten different temperatures (°C). The data is also shown Figure EV 2, but here with k-means clustering. The six charts show the individual clusters from this analysis. From left to right, the clusters were centered around different temperatures (low to high).

**A,** The charts show strains that were categorized as "switch-like", which means that they typically switched from fast growth to no growth within a small temperature range.

**B,** The charts show TS mutants categorized as "gradually" switching.

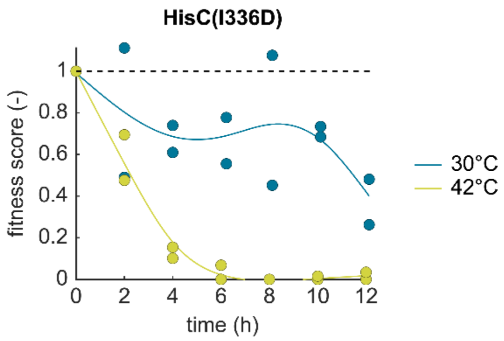


**Supplementary Figure 6 – Fitness score dynamics of the TS mutant $HisC^{I336D}$ during the growth competition experiment.**

Dots indicate n=2 replicates. Blue color indicates data from the 30°C culture, and yellow shows data from the 42°C culture.
